# Nutrient Enrichment and Connectivity Jointly Shape Bacterioplankton Diversity

**DOI:** 10.1101/2025.06.20.660555

**Authors:** Akhil Kholwadwala, Egor Katkov, Patrick Lypaczewski, Andrew Gonzalez, Rowan D.H. Barrett, B. Jesse Shapiro

## Abstract

It is increasingly important to understand the response of freshwater communities and ecosystems to fertilizers given their widespread usage and the propensity for these fertilizers to runoff into rivers and lakes. Dispersal, an important ecological factor mediated by landscape connectivity, could potentially counteract the impacts of anthropogenic stressors through the reintroduction of communities unperturbed by local stressors. However, this potential has not yet been studied in the context of nutrient stressed natural communities. Here, we investigate the impacts of nutrient enrichment and connectivity on freshwater bacterioplankton communities. We subjected mesocosms stocked with native bacterioplankton communities to different combinations of nutrient enrichment and connectivity (volumes of water transferred between mesocosms). We show that nutrient enrichment strongly structures the bacterioplankton community, favoring nutrient tolerant taxa and depressing taxonomic diversity. Connectivity, however, interacts with nutrient enrichment to restore functional diversity in communities subjected to the highest levels of nutrient stress. Despite the ameliorating effects of dispersal, nutrient enrichment leaves a consistent signature in communities, driving a shift from more heterotrophic to more phototrophic communities. Taken together, our results demonstrate that while nutrient enrichment significantly impacts freshwater bacterioplankton communities, connectivity can help restore functional diversity to a certain extent.

## 1. Introduction

Freshwater ecosystems around the world are faced with anthropogenic change, notably from pesticides and fertilizers, changes in land use and water management, the spread of invasive species and diseases, and the growing effects of climate change [1,2]. The individual and more-often combined impacts of these stressors have consequential effects with estimates suggesting that up to a quarter of freshwater animal species are threatened with extinction [2]. This extinction risk is demonstrative of other trends, with these stressors driving shifts in community composition and declines in overall richness and biodiversity across freshwater taxa [3]. Depending on the levels of functional redundancy within the community, the local extinctions and shifts in community composition could have significant ramifications, altering ecosystem function and potentially driving declines in ecosystem productivity and services [4].

Given the magnitude of the threats and the clear consequences for species, communities, and ecosystems, it is of increasing interest to determine if and how communities facing persistent stressors can recover. While “community rescue” — by which community members adapt or acclimate to the stressor and thereby recover — is possible [5,6], a plausible alternative mechanism, particularly in more interconnected freshwater ecosystems, involves dispersal. In freshwater ecosystems, connectivity (the displacement of water and organisms) is an important force maintaining biodiversity [3]. Consequently, connectivity-mediated dispersal, particularly from unexposed communities, could allow stress-exposed communities to maintain higher diversity under continuous exposure to the stressor(s), allowing for spatial insurance [7], thereby preserving ecosystem function. By introducing species from unexposed communities, dispersal and connectivity can serve as a counteracting force to the changes in species composition caused by local stressors [8]. Further, over long timescales, connectivity allows species dispersal which provides access to the regional pool of species and can allow disturbed communities to recover upon the abatement of the stressor [9].

Nutrient (both nitrogen and phosphorous) enrichment from agricultural runoff and urban discharge represents a major threat to freshwater ecosystems and biodiversity [10,11]. Unfortunately, this threat is only projected to grow; a total of 187.95 megatons of fertilizers, including 109.19 megatons of nitrogen-based fertilizer and 44.11 megatons of phosphorus-based fertilizer, were used across the globe in 2022 alone [12]. This usage is projected to set a record high in 2025 with a forecasted total of 205 megatons of fertilizers [13]. In addition, due to land surface changes and changes in precipitation and potential evapotranspiration, runoff is projected to increase globally through the end of the century [14].

Bacterioplankton play important roles in these aquatic ecosystems by recycling organic matter and nutrients back into the ecosystem in a process known as the microbial loop [15]. Anthropogenic disruptions to bacterioplankton communities, depending on the nature and persistence of the stressor, often alter overall community function, including processes like nutrient cycling [16]. Therefore, evaluating bacterioplankton dynamics in response to nutrient enrichment is essential in any discussion of the health of freshwater ecosystems.

Previous studies have indicated that nutrient enrichment and the resulting eutrophication drive changes in bacterioplankton community composition, likely due to deterministic processes such as species sorting – filtering driven by the local environmental and ecological conditions [17–19]. A well-documented, often extreme, example of the effects of nitrogen and phosphorus enrichment on microbial community composition are cyanobacterial or algal blooms [20,21]. Beyond the increased abundance of cyanobacterial species, these blooms have broader impacts on bacterioplankton communities and their composition, altering biotic interactions and the abiotic environment, particularly as it relates to the availability and composition of dissolved organic matter [22,23]. In the context of nutrient enrichment and eutrophication, dispersal has primarily been considered a stochastic process that adds uncertainly to the otherwise deterministic process of species sorting [18]. However, we propose that dispersal could also serve as a source of resistance (defined in this context as maintaining a community composition more similar to the pristine state) and play a more deterministic role by counteracting the nutrient-induced changes in community composition.

To test the potential for connectivity to drive community resistance in the face of an anthropogenic stressor, we designed a factorial experiment to evaluate the response of freshwater bacterioplankton communities to different levels of nutrient enrichment and connectivity. We conducted this experiment in mesocosms at McGill University’s Large Experimental Array of Ponds (LEAP), stocked with bacterioplankton communities from a pristine mesotrophic hilltop lake. We hypothesized that (i) nutrient enrichment strongly structures bacterioplankton communities by driving declines in taxonomic and functional diversity, but that (ii) high levels of connectivity allow community resistance to nutrient-induced changes, maintaining higher taxonomic and functional diversity in the face of nutrient stress.

## 2. Materials and Methods

### (a) Experimental design and sampling

We conducted this study at McGill University’s Large Experimental Array of Ponds (LEAP) located at the Gault Nature Reserve at Mont Saint Hilaire, Québec, Canada (45°32’06.3’N 73°08’54.5’W). Each of the 96 1,136-liter mesocosms (Rubbermaid) in this array is supplied with freshwater and native planktonic communities from the nearby Lac Hertel, a hilltop lake supplied only by rainwater with no known history of agricultural contamination. Our study applied nine combinations of three different levels of connectivity (0, 20, and 40 % of volume transferred/week) and nutrient addition (0, 50, and 200 µg of phosphorus L^-1^ week^-1^) to 17 mesocosms to evaluate the impacts of connectivity and nutrient enrichment on overall bacterioplankton community dynamics (Fig. 1).

**Figure 1:**
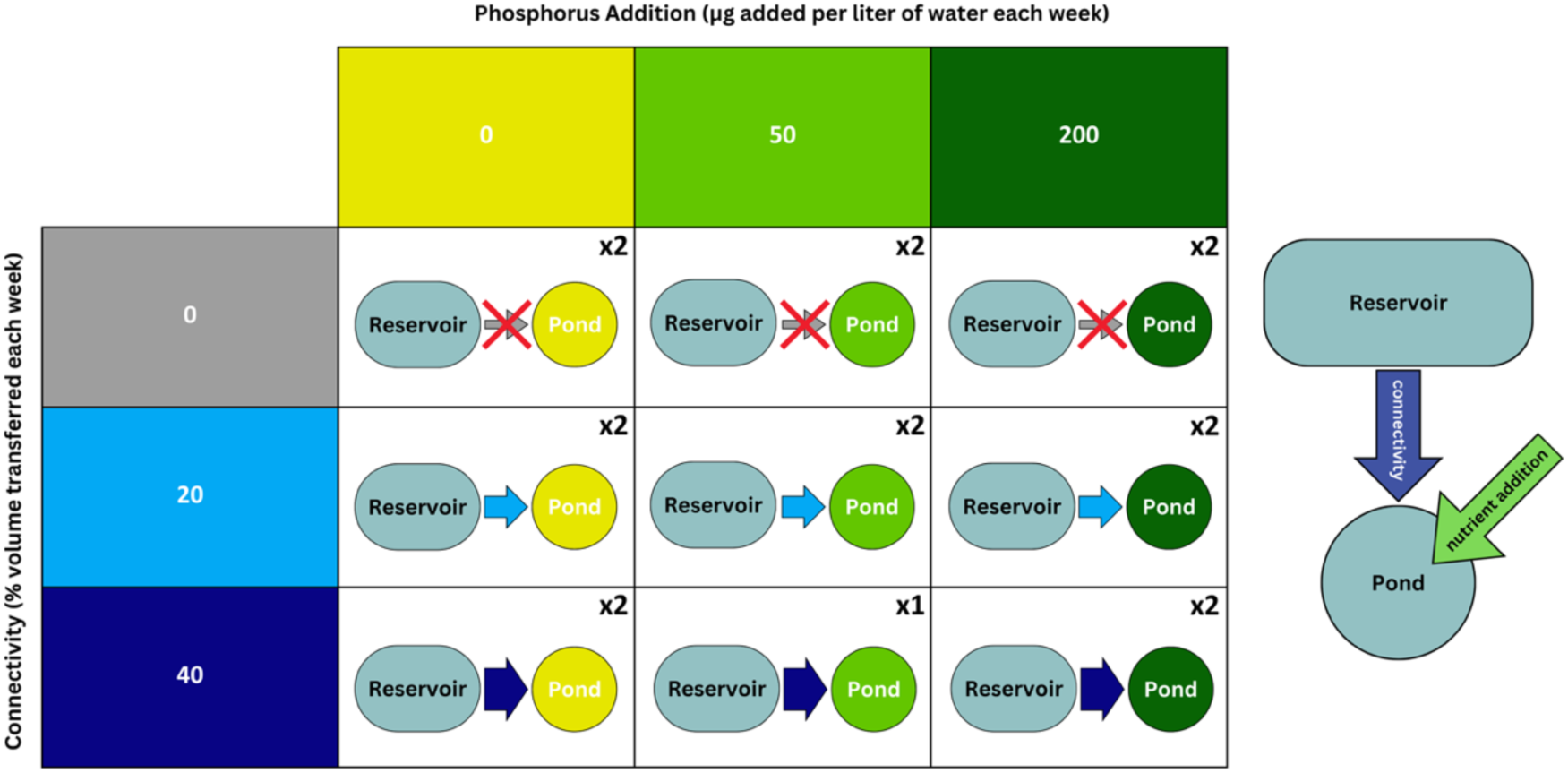
Factorial experimental design to test nutrient stress, connectivity, and their potential interaction. Mesocosms were subjected to nine different combinations of nutrient addition and connectivity. Note that the color key (shades of grey to blue for connectivity, shades of yellow to green for nutrients) is consistent in subsequent figures.

We cleaned the mesocosms, filled them with lake water held in a reservoir, and allowed them to acclimate for a period of one week, before subjecting them to weekly connectivity and nutrient treatments over 15 weeks from July 4^th^ to October 13^th^, 2022. The connectivity treatment was applied on a weekly basis; a 10 L graduated bucket was used to first remove the appropriate volume from the mesocosm before replacing it with water from the reservoir. To control for the turbulence provided by this process, the bucket was also used to remove and add back in the same water from mesocosms that received less than the maximum 40% connectivity treatment. The weekly nutrient treatment directly followed the connectivity treatment, and phosphorous (both KH_2_PO_4_ and K_2_HPO_4_) and nitrogen (KNO_3_) were added in a molar ratio of 1:31 to maintain the P:N molar ratio found in Lac Hertel [24]. These treatments were applied to 17 mesocosms, including two physical replicates for all but one of the 9 treatment combinations.

We collected samples at two timepoints during the experiment on August 18^th^, 2022, and October 4^th^, 2022, giving us two temporal replicates for each of our ponds. For each sample, a PVC pipe with a one-inch diameter was used to sample 1 L of the community and capture diversity across the vertical environmental gradient present in water bodies. The 1 L sample was homogenized, and 200 mL was vacuum filtered through a 0.22-micron PES membrane filter (Millipore). 200 mL of Millipore water was also filtered as filtration negative control to account for contamination. All sample filters were subsequently placed into 2 mL microcentrifuge tubes, placed on dry ice for transport, and stored at -80 °C until extraction.

### (b) DNA extraction, sequencing, and classification

For DNA extraction, filters were cut in half; one half was returned to the sample tube and stored back at -80 °C while the other half was extracted following a modified protocol with the Qiagen DNeasy® PowerWater® Kit. As an extraction negative control, a new, unused filter was extracted using the same process. The extracted DNA was quantified using a Qubit Fluorometer (ThermoFisher) and the Qubit High Sensitivity dsDNA assay (ThermoFisher).

Following quantification, 36 different 27F forward primers (electronic supplementary material, Table S1) and the 1492R reverse primer (5’-CGGYTACCTTGTTACGACTT-3’) were used to amplify and multiplex the entire length of the bacterial 16S rRNA gene [25,26]. For further multiplexing, two pairs of Nanopore native barcodes were ligated onto the amplified DNA sequences (electronic supplementary material, Table S2). The samples were subsequently sequenced using the PromethION P2 from Oxford Nanopore Technologies (ONT). The resulting output files were basecalled and demultiplexed using the super-accurate basecalling model and demultiplexed using Guppy v6.5.7 (ONT) with the dna_r10.4.1_e8.2_400bps_sup model and a minimum Q-score cut off of 7 (the custom barcoding arrangement file and custom barcodes file have been made publicly available [27]) [28].

Sequences were subsequently imported into Linux (Ubuntu 22.04.5) for taxonomic and functional assignment. Given the potentially higher error rates of long-read sequencing technologies such as ONT compared to Illumina, we opted to use Emu, a classifier designed for long-read 16S sequences, for taxonomic assignment [29,30]. Emu utilizes an expectation-maximization algorithm to iteratively describe and update the taxonomic profile of the sample based on updating probabilities of taxonomic assignments for each read. Using an abundance threshold of 0.001, we were able to generate taxonomic profiles for all of our samples. In total, prior to filtering, we had a total of 17 764 551 sequence reads classified down to 938 species-level classifications with an average read depth of 510 007 (SD: 367 258) across our 34 samples compared to an average read depth of 106 079 (SD: 154 658) across our four negative controls. We also included a mock community (MSA-1002^TM^, ATCC) made up of a 20-strain even mix as a positive control. We subsequently utilized FAPROTAX [31] which maps taxonomic assignments to functional categories, to generate estimates of functional diversity.

### (c) Statistical analysis

Statistical analysis was conducted in RStudio with R version 4.4.0 (2024-04-24) using packages readxl [32], ggplot2 [33], dplyr [34], BiocManager [35], phyloseq [36], decontam [37], ANCOMBC [38,39], FSA [40], vegan [41], tidyr [42], forcats [43], and tidypaleo [44]. Beginning with our taxonomic analysis, the output from Emu was first converted into a phyloseq object. We subsequently used the R-package decontam [37] to statistically identify and remove contaminating taxa based on their high abundance in the negative controls. Out of 938 taxa, decontam identified 10 as contaminants, and they were subsequently removed from our data set (electronic supplementary material, Table S3). We further filtered our data to only retain taxa present in our 34 samples leaving a total of 731 species.

To evaluate the effects of our treatments on alpha diversity, we first calculated both the species richness and Shannon index for each of the samples, using the Kruskal-Wallis test and Dunn test to test for any significant differences between the levels within each of our treatment types. We subsequently calculated both the Jaccard distance and Bray-Curtis distance, using permanova and beta-dispersion tests to evaluate the effects of the nutrient and connectivity treatments on beta diversity, testing, respectively, for significant differences in the centroids and the dispersions when the samples were grouped according to treatment. We used ANCOMBC2 for our differential abundance analysis as it detects differences in relative abundance after correcting for compositional effects [38,39]. We identified differentially abundant taxa between nutrient enriched (50 µg/L/week and 200 µg/L/week) and control (0 µg/L/week) samples at three taxonomic levels (phylum, class, and order) using a prevalence threshold of 0.1 and adjusting p-values with the Holm method. For a more nuanced analysis, we constructed generalized linear models (GLM) to look at alpha-diversity as a function of the individual fixed and combined interaction effects of nutrient enrichment and connectivity level. We added an additional fixed effect of sampling date to account for the fact that our data consists of samples taken on two separate dates nearly 7 weeks apart. We fit our model for species richness to a Poisson distribution, used for count data, and that for Shannon diversity to a Gaussian distribution, for normally distributed data.

For the analysis of the functional diversity, we first identified functional classes that had different abundances between nutrient enriched (50 µg/L/week and 200 µg/L/week) and control (0 µg/L/week) samples. We filtered our data to only include functional classes present in at least 10% of the samples before evaluating statistically significant differences in abundance with the Mann-Whitney U-test and correcting for multiple testing using the Holm method. To understand how the overall functional diversity changes as a function of our two treatments, we constructed a GLM testing the number of functional classes present (functional richness) as a function of the fixed effects of nutrient enrichment, connectivity level, and sampling date and an interaction effect between our two treatments. We fit this model to a Poisson distribution.

## 3. Results

### (a) Nutrient addition reduces diversity and changes community structure

To test how nutrients and connectivity jointly affect bacterioplankton communities, we analyzed 34 samples taken across two timepoints from mesocosms subjected to different combinations of nutrient enrichment and connectivity. We used long-read 16S rRNA amplicon sequencing (see Methods) to quantify the relative abundances of prokaryotic taxa within communities.

We first evaluated the impacts of connectivity and nutrient enrichment on community alpha diversity. We found a significant effect of nutrient treatment reducing species richness at both levels of 50 µg/L/week (Z = 2.63, holm-adjusted *p* = 0.017) and 200 µg/L/week (Z = 3.57, holm-adjusted *p* = 0.0011) (Fig. 2). Similarly, nutrient treatment reduced Shannon diversity at both tested concentrations (Z = 3.48, holm-adjusted *p* < 0.001 and Z = 3.75, holm-adjusted *p* < 0.001, respectively) when compared to the control samples (Fig. 2). In contrast, connectivity alone had no significant impact on the alpha-diversity of the bacterioplankton communities (Fig. 2; holm-adjusted *p*-values all > 0.05).

**Figure 2:**
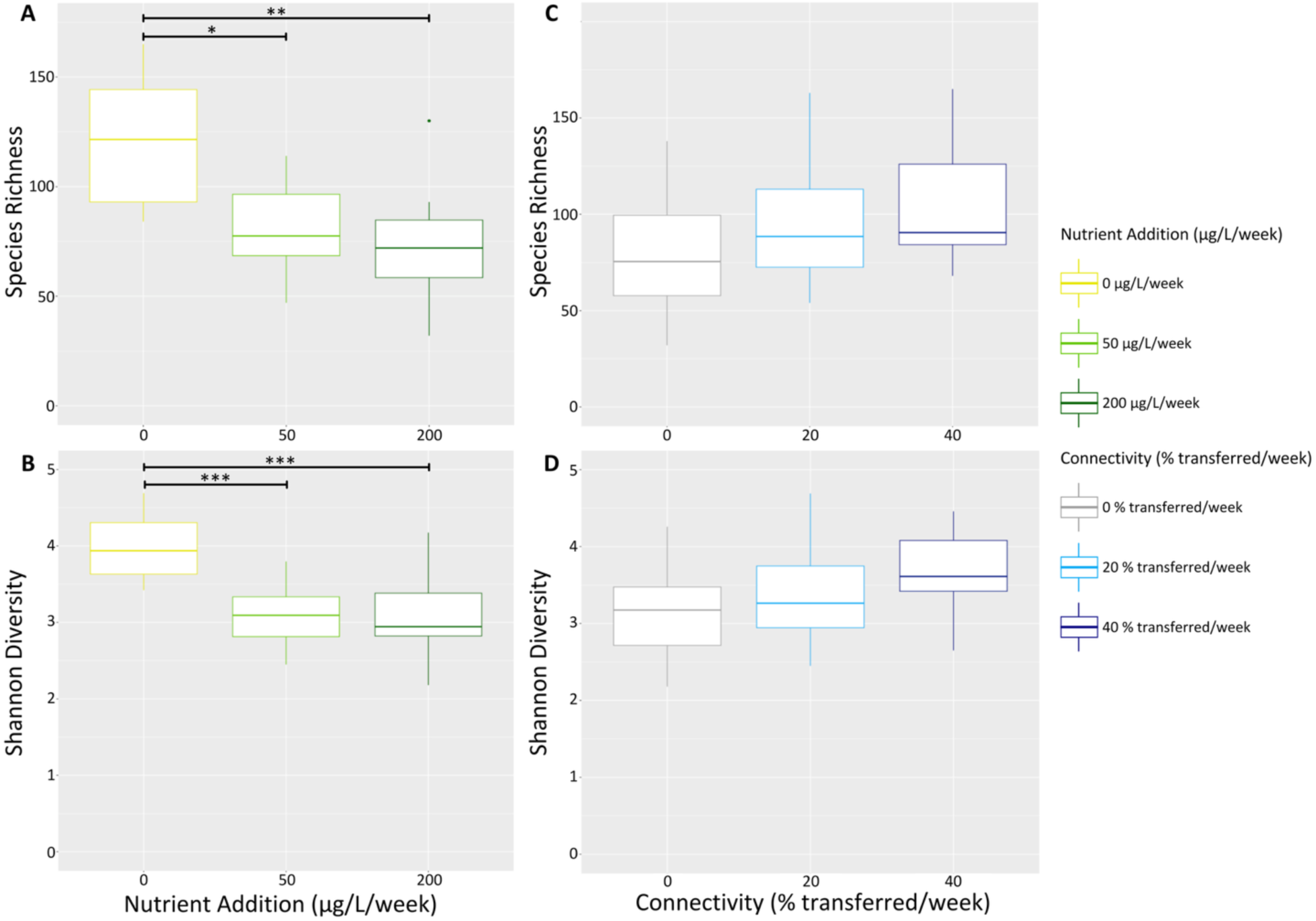
Nutrient addition significantly reduces community diversity, but connectivity has no significant effect. Box plots showing the independent effects of nutrient enrichment and connectivity on two measures of alpha-diversity (species richness and Shannon diversity). Boxes show the interquartile range, with the horizontal line within the box showing the median and whiskers showing the spread of the values, excluding outliers. Asterisks indicate significant differences after multiple testing correction: (*) p < 0.05, (**) p < 0.01, (***) p < 0.001

Next, we asked whether nutrient-induced losses in diversity were accompanied by changes in community composition. Two measures of differences in community composition (beta diversity) were significantly different across nutrient treatment groups (Fig. 3). The three nutrient treatments had significantly different centroids (Jaccard distance: F_2,31_ = 3.213, *p* = 0.001, Bray-Curtis distance: F_2,31_ = 5.096, *p* = 0.001) but not significantly different dispersions (Jaccard distance: F_2,31_ = 0.187, *p* = 0.830, Bray-Curtis distance: F_2,31_ = 0.193, *p* = 0.826). These effects were driven by both levels of nutrient additions having distinct community compositions relative to control communities (Fig. 3A, C). However, neither measure of beta-diversity was significantly different between the connectivity levels (Fig. 3B, D), with no significant difference in the centroids (Jaccard distance: F_2,31_ = 1.048, *p* = 0.288, Bray-Curtis distance: F_2,31_ = 1.054, *p* = 0.308) or the dispersions of the different connectivity groups (Jaccard distance: F_2,31_ = 0.572, *p* = 0.578, Bray-Curtis distance: F_2,31_ = 0.230, *p* = 0.795). Further, nutrient enrichment alone explains 17.2% and 24.7% of the variation in community structure, when considering Jaccard distance and Bray-Curtis distance respectively. By contrast, connectivity alone explained only 6.3% and 6.4%.

**Figure 3:**
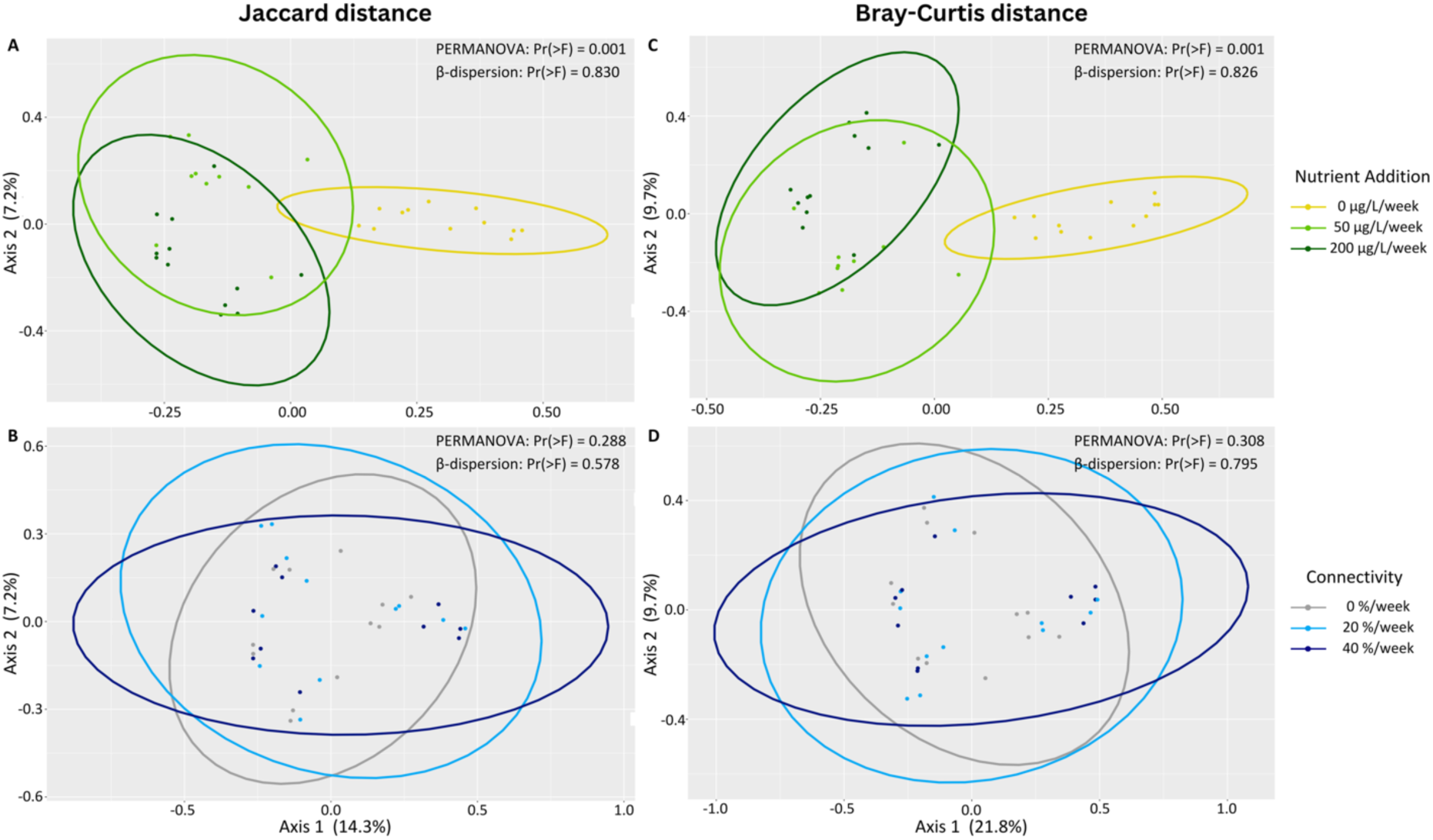
Nutrient addition, but not connectivity, significantly changes community composition. PCoA plots of our samples based on Jaccard distance (A and B) and Bray-Curtis distance (C and D). The results of the PERMANOVA to test the for significant differences between the centroids and the beta-dispersion test to test for significant differences in the dispersions of the groups are displayed.

These nutrient-induced changes in community composition are due to changes in the relative abundance of different phyla, classes, and orders within the community (electronic supplementary material, Fig. S1). To identify taxa driving the changes in beta diversity, we compared the relative abundances between control samples (0 µg/L/week) and nutrient enriched samples (50 µg/L/week and 200 µg/L/week combined). At the phylum level, *Actinobacteria* is relatively more abundant in control ponds (*W* = 9.73, holm-adjusted *p* < 0.001) while *Bacteroidetes* is enriched in nutrient-treated ponds (*W* = -3.20, holm-adjusted *p* = 0.030) (electronic supplementary material, Fig. S2). The association between nutrient treatment and *Bacteroidetes* is driven by increased abundances of both *Sphingobacteriales* and *Cytophagales* (*W* = -6.01, holm-adjusted *p* < 0.001) at the order level.

Within *Proteobacteria,* the classes *Gammaproteobacteria* (*W* = -4.16, holm-adjusted *p* = 0.0077) and *Alphaproteobacteria* demonstrate opposite trends to *Betaproteobacteria* (*W* = 4.91, holm-adjusted *p* < 0.001), with the former more abundant in enriched samples and the latter more abundant in control samples (electronic supplementary material, Fig. S2). Taxa within *Gammaproteobacteria* and *Betaproteobacteria* show consistent responses with the order *Xanthomonadales* (*W* = -4.90, holm-adjusted *p* = 0.0035) driving the trends in the former and the orders *Nitrosomonadales* and *Burkholderiales* (*W* = 3.89, holm-adjusted *p* = 0.029) responsible for the trends observed in the latter. On the other hand, taxonomic orders within *Alphaproteobacteria* have varied responses to enrichment with *Pelagibacterales* more abundant in control samples (*W* = 9.27, holm-adjusted *p* = 0.0081) and *Rhodospirillales* more abundant in nutrient enriched samples (*W* = -4.31, holm-adjusted *p* = 0.0058).

At the order level, we find the several *Planctomycetes* orders enriched in control samples (*W* = 5.56, holm-adjusted *p* = 0.027, *W* = 5.52, holm-adjusted *p* = 0.027) (electronic supplementary material, Fig. S2). On the other hand, we find the cyanobacterium *Chroococcales* to be enriched in nutrient-treated ponds relative to controls (*W* = -4.23, holm-adjusted *p* = 0.041)

### (b) Interactive effects of nutrients and connectivity on community diversity

For a more nuanced evaluation of the combined effects of nutrient enrichment and connectivity, we constructed generalized linear models (GLMs) describing alpha-diversity (either species richness or Shannon diversity) as a function of the fixed effects of nutrient enrichment, connectivity level, and sampling date, and an interaction effect between nutrient enrichment and connectivity (Fig. 4 and electronic supplementary material, Table S4). The only significant effect in the Shannon diversity model is nutrient enrichment (estimate = - 4.66e-3, *t* = -3.11, *p* = 0.0042). The model for species richness meanwhile shows a significant negative effect of nutrient enrichment (estimate = -3.73e-3, *Z* = -9.86, *p* < 0.001) and of sampling date (estimate = -2.64e-1, *Z* = -7.36, *p* < 0.001). These effects are consistent with our univariate analyses (Fig. 2) and also reveal a slight decline in diversity over time. However, GLMs also identified a significant positive effect of connectivity level on richness (estimate = 2.83e-3, *Z* = 1.96, *p* = 0.050), suggesting a subtle effect of connectivity restoring diversity. We also found a significant interaction between nutrient enrichment and connectivity (estimate = 6.06e-5, *Z* = 4.48, *p* < 0.001) on species richness, with connectivity driving higher species richness at higher levels of nutrient enrichment (Fig. 4A). Although not significant in the model, a similar effect is visually evident for Shannon diversity as well (Fig. 4B). Together, these results suggest that, at the highest levels of nutrient stress, connectivity could play a role in restoring diversity.

**Figure 4:**
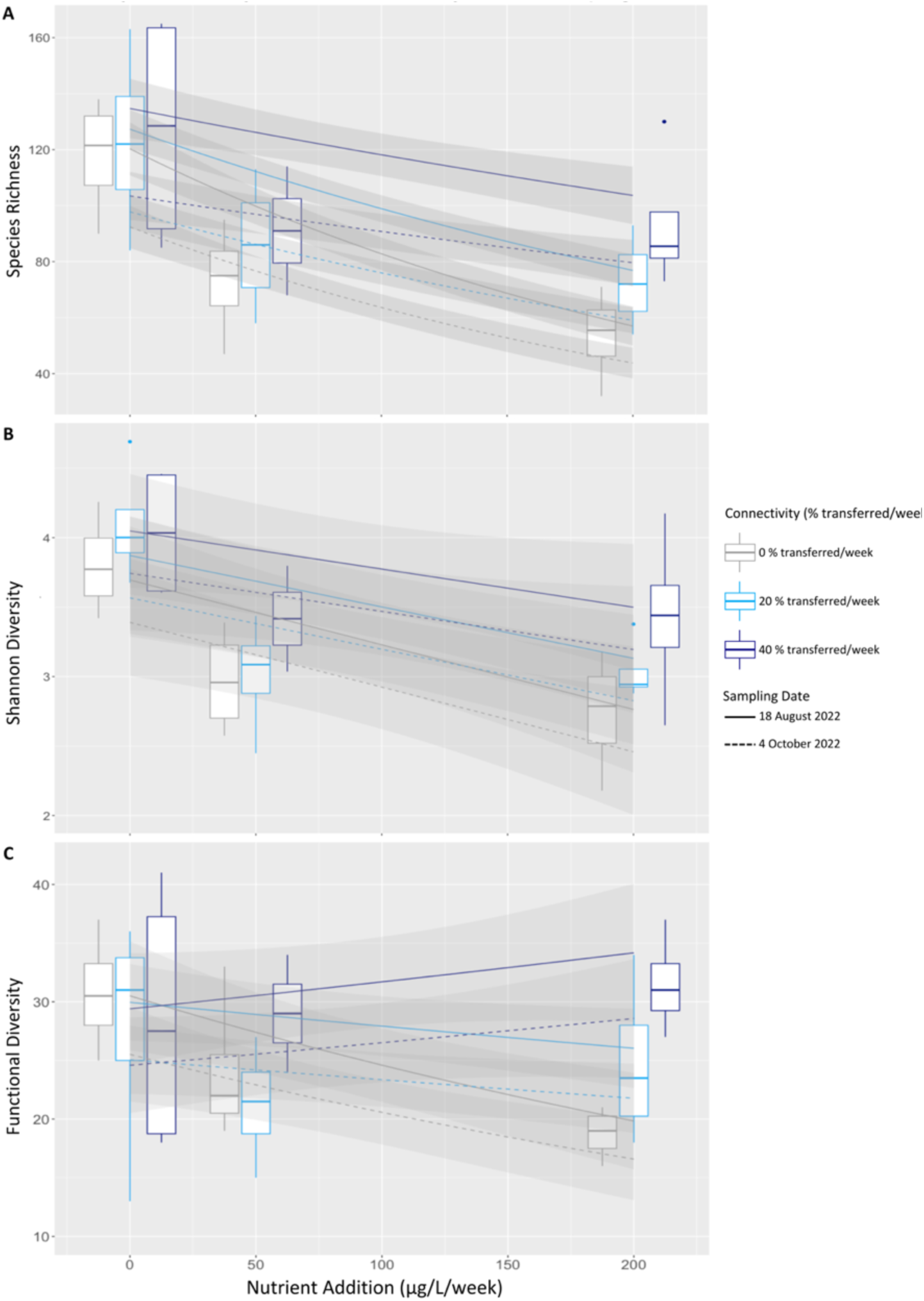
Connectivity restores taxonomic and functional diversity at high levels of nutrient stress. A box plot showing the (A) observed species richness, (B) Shannon diversity, and (C) functional diversity across all combinations of nutrient addition and connectivity. The plot is overlayed with a generalized linear mixed model describing diversity as a function of nutrient addition, connectivity level, and sampling date. The shaded region defines the 95% confidence interval.

### (c) Nutrient addition and connectivity shape the functional composition and diversity of bacterioplankton communities

To evaluate the broader–and potentially ecosystem-level–impacts of the changes in bacterioplankton community composition and alpha diversity, we analyzed the impacts of nutrient enrichment and connectivity on the inferred functional diversity of taxa identified in our ponds. The 731 species-level taxa across our samples and treatment combinations were successfully mapped to 61 non-exclusive functional categories (see Methods).

Given the strength of the nutrient enrichment in structuring bacterioplankton community composition and alpha diversity (Figs. 2, 3, 4), we evaluated how nutrient enrichment alone shapes the relative abundances of the various functional classes attributed to the taxa present in the sampled communities. Specifically, we compared functional classes whose relative abundances differ between the enriched (50 µg/L/week and 200 µg/L/week) and control samples (0 µg/L/week). We found that phototrophy was associated with nutrient enrichment (*W* = 245, holm-adjusted *p* < 0.001), particularly driven by an increase in photoheterotrophy in nutrient enriched ponds (*W* = 229, holm-adjusted *p* = 0.0085) (electronic supplementary material, Fig. S3).

In contrast, overall chemoheterotrophy declined in nutrient enriched samples, although the difference was not significant after correcting for multiple tests (electronic supplementary material, Fig. S3). Several heterotrophic functional classes relating to the degradation of polysaccharides declined in nutrient enriched samples, including xylanolysis (*W* = 37.5, holm-adjusted *p* = 0.0028) and cellulolysis (*W* = 69, holm-adjusted *p* = 0.052). We also observed a decline in nitrogen fixation (*W* = 44, holm-adjusted *p* = 0.050) in nutrient enriched samples.

Finally, we used a GLM to test how the fixed effects of nutrient enrichment, connectivity level, their interaction, and time of sampling affected the inferred number of functional classes (Fig. 4C and electronic supplementary material, Table S5). In addition to significant negative effects of nutrient enrichment (estimate = -2.15e-3, *Z* = -3.25 *p* = 0.0012) and sampling date (estimate = -1.79e-1, *Z* = -2.65, *p* = 0.0081), the GLM shows a significant positive interaction effect between connectivity and nutrient enrichment (estimate = 7.26e-5, *Z* = 3.00, *p* = 0.0027), with higher levels of connectivity maintaining higher functional richness in samples as the level of nutrient enrichment increases. This parallels the effect observed for species richness (Fig. 4A) and suggests that connectivity could promote functional resistance in the face of nutrient stress.

## 4. Discussion

### (a) Nutrient enrichment strongly structures bacterioplankton communities

As expected, our results demonstrate that nutrient enrichment, even at intermediate levels, strongly structures the bacterioplankton community composition. Consistent with prior work, our analysis suggests that the long-term effects of nutrient enrichment on community composition are selective. The directed shifts in community composition following nutrient addition confirm the predominant effects of species sorting–particularly in response to nutrients and eutrophication–in shaping aquatic bacterial community composition [17,18,45]. Given that the transfer of water to all mesocosms occurred from a common reservoir, the lack of homogenization of the bacterioplankton community composition at the highest level of connectivity suggests that the sorting effects due to enrichment overcome any mass effects, in which high levels of dispersal or connectivity maintain taxa not well adapted to the local environmental conditions [45].

The evidence for nutrient enrichment driving a species sorting process that shapes bacterioplankton community composition is furthered bolstered when analyzing the taxa whose abundances are driving the differences in community composition between the nutrient enriched and control samples. Nutrient tolerant phyla such as certain *Cyanobacteria* (particularly in the order *Chroococcales*) have significantly higher relative abundance in nutrient enriched samples [20]. Taxa within the phylum *Bacteroidetes*, particularly *Sphingobacteriales* and *Cytophagales*, also show increased relative abundance under nutrient enrichment. Members of *Sphingobacteriales* demonstrate a strong chemotactic response to inorganic nutrients like nitrogen and phosphorous [46,47] and the ability to lyse cyanobacterial cells [48]. Further, members of the *Bacteroidetes* and *Alphaproteobacteria,* including *Rhodospirillales,* have been associated with the phycosphere of cyanobacteria, potentially playing an important role in algal growth and likely making them more responsive to nutrient addition and any accompanying cyanobacterial bloom [49,50]. The order *Gammaproteobacteria* is also more abundant in nutrient enriched samples, which is consistent with descriptions of their rapid growth in response to nutrient addition, potentially allowing them to outcompete other taxa [51].

In contrast, taxa that are generally regarded as oligotrophs (better competitors in low nutrient conditions) such as *Pelagibacterales* [52,53], *Verrucomicrobia* [54], and *Actinobacteria* [48,51,55] show higher relative abundance in the control conditions, as expected. However, the described life strategies of *Verrucomicrobia* do appear to vary between studies with others describing the taxa as more abundant in more eutrophic environments [56,57]. Similarly, the trends observed in our experiment for *Planctomycetes* and *Betaproteobacteria*, which have higher relative abundance in our control samples, contradict previous studies which describe correlations between *Planctomycetes* abundance and algal blooms [58] and associations between *Betaproteobacteria* and cyanobacteria [51] and increased abundance of *Betaproteobacteria* in nutrient enriched environments [17].

Shifts in the community composition are also accompanied, at a functional level, by declines in the relative abundance of nitrogen fixing functional classes in our nutrient enriched samples. With increased concentrations of bioavailable nitrogen (NO_3_) from nutrient enrichment, bacterial fixation of atmospheric nitrogen could be selected against. Together, these results support the notion of a deterministic, selective shift in the bacterioplankton communities following nutrient addition at concentrations consistent with recorded nutrient concentrations in lakes across the globe [59].

### (b) Nutrient enrichment and connectivity interact to shape alpha-diversity

Nutrient enrichment has strong impacts on community alpha diversity, driving a decline in richness and Shannon diversity with increasing enrichment, with a 41% decline in the median species richness in the most nutrient enriched samples compared to the unenriched ones. While some species loss could be stochastic, our taxonomic results instead suggest that the nutrient treatment effectively filters out species unable to tolerate or acclimate to enriched conditions, thereby reducing overall diversity. This effect is most notable when connectivity is absent. As per our hypothesis, connectivity does, to some extent, maintain alpha diversity. Further, as indicated by the significant interaction effects between the two treatments, connectivity has a stronger effect in increasing alpha diversity in communities subject to the highest levels of nutrient enrichment, with high levels of connectivity boosting the median species richness by 54% under the highest levels of nutrient enrichment compared to just 6% in unenriched communities. The taxonomic diversity of communities exposed to any level of nutrient treatment, however, does remain significantly lower than control communities, suggesting, perhaps, that connectivity alone cannot successfully rescue stressed communities under these conditions.

### (c) Connectivity sustains functional diversity

Given evidence of functional redundancy in bacterioplankton, including from previous research conducted at our study site [60], ecosystem function may be maintained following disturbance and declines in alpha diversity. Our results indicate a decline in functional diversity with increasing nutrient enrichment, but because taxonomic richness declines more rapidly than functional richness as a function of nutrient enrichment, there is also evidence of functional redundancy within the bacterioplankton community in our study. In addition, the functional diversity results indicate an outsized role of connectivity maintaining overall functional diversity in the community, and therefore, likely, overall ecosystem function as well, irrespective of declines in taxonomic diversity. While communities see a 33 % decline in taxonomic richness under high levels of connectivity in nutrient enriched environments, high levels of connectivity maintain the functional diversity within nutrient enriched communities relative to controls.

While connectivity may not substantially increase the taxonomic richness enough to restore overall community diversity, high levels of connectivity can maintain the functional diversity of stressed communities. This suggests that, while the magnitude of the stressor prevents the successful colonization of the majority of dispersing bacteria, tolerant taxa with diverse functions are successfully able to establish themselves given sustained bouts of connectivity in heterogenous landscapes, maintaining ecosystem function and productivity in line with the spatial insurance hypothesis [7].

### (d) Nutrient enrichment drives a heterotrophy to phototrophy shift

While functional diversity appears to have been maintained by high levels of connectivity, there appears to be a broad regime shift in the community from heterotrophy to phototrophy under nutrient enrichment. Control communities show higher abundances of chemoheterotrophic functional classes whereas communities subjected to nutrient enrichment show significantly higher abundances of phototrophic classes. Notably, this trend is dominated by increased abundance of *Cyanobacteria* of which many taxa are described as photoheterotrophic [61]. This shift is indicative of eutrophication and undoubtedly impacts the functioning of the microbial loop, with reductions in carbon recycling and nutrient remineralization [62]. The subsequent impacts are likely limited in our experimental system, with its continuous input of inorganic nutrients. However, the impacts would be highly consequential in systems with low connectivity in which the nutrient enrichment occurs seasonally or intermittently, thereby leaving long stretches of time without an exterior supply of inorganic nutrients and insufficient remineralization of nutrients by heterotrophic bacteria to sustain local communities.

### (e) Caveats

There are several important caveats to our study. First, as our samples were taken 6 and 13 weeks after the start of the experiment, our results capture variation within summer and fall seasons, but not potentially rapid daily dynamics, nor longer-term effects. In addition, the nature of the study design means that the connectivity treatment simultaneously (re)introduces new taxa and dilutes the nutrient stress. Consequently, it is difficult to ascertain whether richness and functional diversity recovers primarily through the abatement of the stressor or the introduction of diverse and nutrient tolerant taxa. This limits our ability to generalize the results to taxa, particularly terrestrial ones, in which dispersal and the magnitude of the stressor are decoupled. However, in the aquatic context it is realistic that stressor concentrations and species migration would be coupled in flow from rivers and streams.

Our 16S rRNA gene amplicon sequence data is somewhat limited in taxonomic resolution, meaning that sub-species (ℌstrain” level) dynamics could differ from what we observed. Moreover, extrapolating from 16S sequences to metabolic functions or lifestyles is imperfect, but we believe that the general categories considered here (*e.g.* phototrophs vs. heterotrophs) are fairly robust. Finally, while our study included weekly nutrient treatments over the entire duration of the experiment, future studies should evaluate the role of connectivity in the rescue and recovery of communities during and following the abatement of the stressor/nutrient treatment.

## 5. Conclusion

Using the experimental mesocosm system at LEAP, we tested the effects of nutrient enrichment on bacterioplankton community composition and diversity, as well as any potential remediating effects of connectivity. Bacterioplankton are crucial for overall ecosystem function, and therefore, investigating the response of bacterioplankton communities to anthropogenic stressors and potential avenues for the recovery of diversity and ecosystem function is essential. Our results confirm an important role for nutrient enrichment in structuring bacterioplankton communities: driving deterministic shifts in favor of nutrient tolerant species at the expense of others, accompanied by general declines in both taxonomic and functional richness. Dispersal, particularly from unstressed environments, has been proposed as a mechanism to “rescue” communities by reintroducing both species lost to the stressor and potentially new species that may have higher stress tolerance. Our study indicates that connectivity has a significant but modest effect in keeping communities resistant to the reduction in taxonomic diversity caused by nutrient stress. However, high levels of connectivity do sustain functional diversity in the most nutrient stressed samples. Despite this, however, the predominant outcome of nutrient enrichment is a regime shift from primarily heterotrophic to autotrophic communities. Taken together these results suggest that certain regimes of connectivity, with increased flow from pristine to polluted water bodies, could help sustain the functional diversity of bacterial communities.

## Data Accessibility

Raw long-read nanopore 16S sequencing reads and accompanying metadata have been deposited in the National Center for Biotechnology Information (NCBI) Sequence Read Archive (SRA) repository (BioProject: PRJNA1277156). The metadata, code, and input files used for the filtering of the data, the statistical analysis, and the generation of the figures have been uploaded to the Dryad Digital Repository [27]

## Authors’ Contributions

A.K.: conceptualization, methodology, software, investigation, formal analysis, visualization, writing—original draft, writing—review and editing. E.K.: conceptualization, methodology, investigation, writing—review and editing. P.L.: methodology, software, writing—review and editing. A.G.: conceptualization, writing—review and editing, funding acquisition. R.D.H.B.: conceptualization, writing—original draft, writing—review and editing, supervision, funding acquisition. B.J.S.: conceptualization, writing—original draft, writing— review and editing, supervision, funding acquisition.

## Acknowledgements

We would like to acknowledge the many people who contributed to the project. We thank Gregor F. Fussman for helping to design and fund the experiment and Michelle Gros and a team of undergraduate students for maintaining and running the experiment. We are also grateful to Emma Derrick and Maxime Guglielmetti for helping to collect samples. Finally, we thank the team at the Gault Nature Reserve for maintaining the infrastructure at the site and providing necessary assistance.

