## Supplemental figures and tables for "Nutrient Enrichment and Connectivity Jointly Shape Bacterioplankton Diversity"

### Supplemental Material

| Barcode # | Sequence | Barcode # | Sequence |
|---|---|---|---|
| HM001 | 5'- AGCAATCGCGCACAGRGTTYGATYMTGGCTCAG -3' | HM019 | 5'- TCTTCCTCCTCAAAGRGTTYGATYMTGGCTCAG -3' |
| HM002 | 5'- AGTGGACCAACAAAGRGTTYGATYMTGGCTCAG -3' | HM020 | 5'- TGCTAATACCAATAGRGTTYGATYMTGGCTCAG -3' |
| HM003 | 5'- ATAATATTCGGCAAGRGTTYGATYMTGGCTCAG -3' | HM021 | 5'- TGGTCTCATGCGTAGRGTTYGATYMTGGCTCAG -3' |
| HM004 | 5'- ATGCTAGACATCGAGRGTTYGATYMTGGCTCAG -3' | HM022 | 5'- TTAAGCACCTGAAAGRGTTYGATYMTGGCTCAG -3' |
| HM005 | 5'- ATGTTAAATGGAGAGRGTTYGATYMTGGCTCAG -3' | HM023 | 5'- TTAGGTGAATTTGAGRGTTYGATYMTGGCTCAG -3' |
| HM006 | 5'- CAATCGATGGGCCAGRGTTYGATYMTGGCTCAG -3' | HM024 | 5'- TTCATAACAGAAGAGRGTTYGATYMTGGCTCAG -3' |
| HM007 | 5'- CAGGAACAACGCCAGRGTTYGATYMTGGCTCAG -3' | HM025 | 5'- TTCATCGGTCTTAAGRGTTYGATYMTGGCTCAG -3' |
| HM008 | 5'- CCGAGATTGGCCGAGRGTTYGATYMTGGCTCAG -3' | HM026 | 5'- AATAGAACTGCCTAGRGTTYGATYMTGGCTCAG -3' |
| HM009 | 5'- CCTCCAACCGCTGAGRGTTYGATYMTGGCTCAG -3' | HM027 | 5'- AATGGCCGGTTCAAGRGTTYGATYMTGGCTCAG -3' |
| HM010 | 5'- CTTGAGGTGAAGAAGRGTTYGATYMTGGCTCAG -3' | HM028 | 5'- ACATTGTTATAGCAGRGTTYGATYMTGGCTCAG -3' |
| HM011 | 5'- GCCCTATAACACAAGRGTTYGATYMTGGCTCAG -3' | HM029 | 5'- ACCGGATCTGCGAAGRGTTYGATYMTGGCTCAG -3' |
| HM012 | 5'- GCGCTCTCCTACGAGRGTTYGATYMTGGCTCAG -3' | HM030 | 5'- ACTCCCAACCAACAGRGTTYGATYMTGGCTCAG -3' |
| HM013 | 5'- GCTTTGCTTGGCGAGRGTTYGATYMTGGCTCAG -3' | HM031 | 5'- ACTTAACGTTAGCAGRGTTYGATYMTGGCTCAG -3' |
| HM014 | 5'- GGCCTATTAAGTTAGRGTTYGATYMTGGCTCAG -3' | HM032 | 5'- AGCCACCGAAGCGAGRGTTYGATYMTGGCTCAG -3' |
| HM015 | 5'- GTATTCGCCTGGTAGRGTTYGATYMTGGCTCAG -3' | HM033 | 5'- AGGCCGTCCTTGTAGRGTTYGATYMTGGCTCAG -3' |
| HM016 | 5'- TAATAGGCTTCTGAGRGTTYGATYMTGGCTCAG -3' | HM034 | 5'- AGTAACTTGTTCCAGRGTTYGATYMTGGCTCAG -3' |
| HM017 | 5'- TATCGCAAGAACAAGRGTTYGATYMTGGCTCAG -3' | HM035 | 5'- CAAGACTCTACACAGRGTTYGATYMTGGCTCAG -3' |
| HM018 | 5'- TCCAGAGAAGAGAAGRGTTYGATYMTGGCTCAG -3' | HM036 | 5'- CAAGTCCACTATGAGRGTTYGATYMTGGCTCAG -3' |

**Table S1:** The forward 27f primers used in conjunction with the 1492R reverse primer (5'-CGGYTACCTTGTTACGACTT-3') to amplify the full-length 16S rRNA gene sequence of prokaryotes in our samples.

| Barcode # | Forward Sequence | Reverse Sequence |
|---|---|---|
| NB01 | 5'- CACAAAGACACCGACAACTTTCTT -3' | 5'- AAGAAAGTTGTCGGTGTCTTTGTG -3' |
| NB02 | 5'- ACAGACGACTACAAACGGAATCGA -3' | 5'- TCGATTCCGTTTGTAGTCGTCTGT -3' |

**Table S2:** The forward and reverse sequences of the nanopore native barcodes used to further multiplex our samples.

| tax_id | species |
|---|---|
| **1082345** | *Sphingobium boeckii* |
| **1147** | *Synechocystis sp. PCC 6714* |
| **1379909** | *Rufibacter sp. DG15C* |
| **1609758** | *Novosphingobium sp. P6W* |
| **1747** | *Cutibacterium acnes* |
| **2518177** | *Flavobacterium sangjuense* |
| **482957** | *Burkholderia lata* |
| **488447** | *Burkholderia contaminans* |
| **653931** | *Sphingomonas alpina* |
| **95486** | *Burkholderia cenocepacia* |

**Table S3:** Taxa identified as contaminants using the R-package decontam (36). These taxa were removed from our samples prior to analysis.

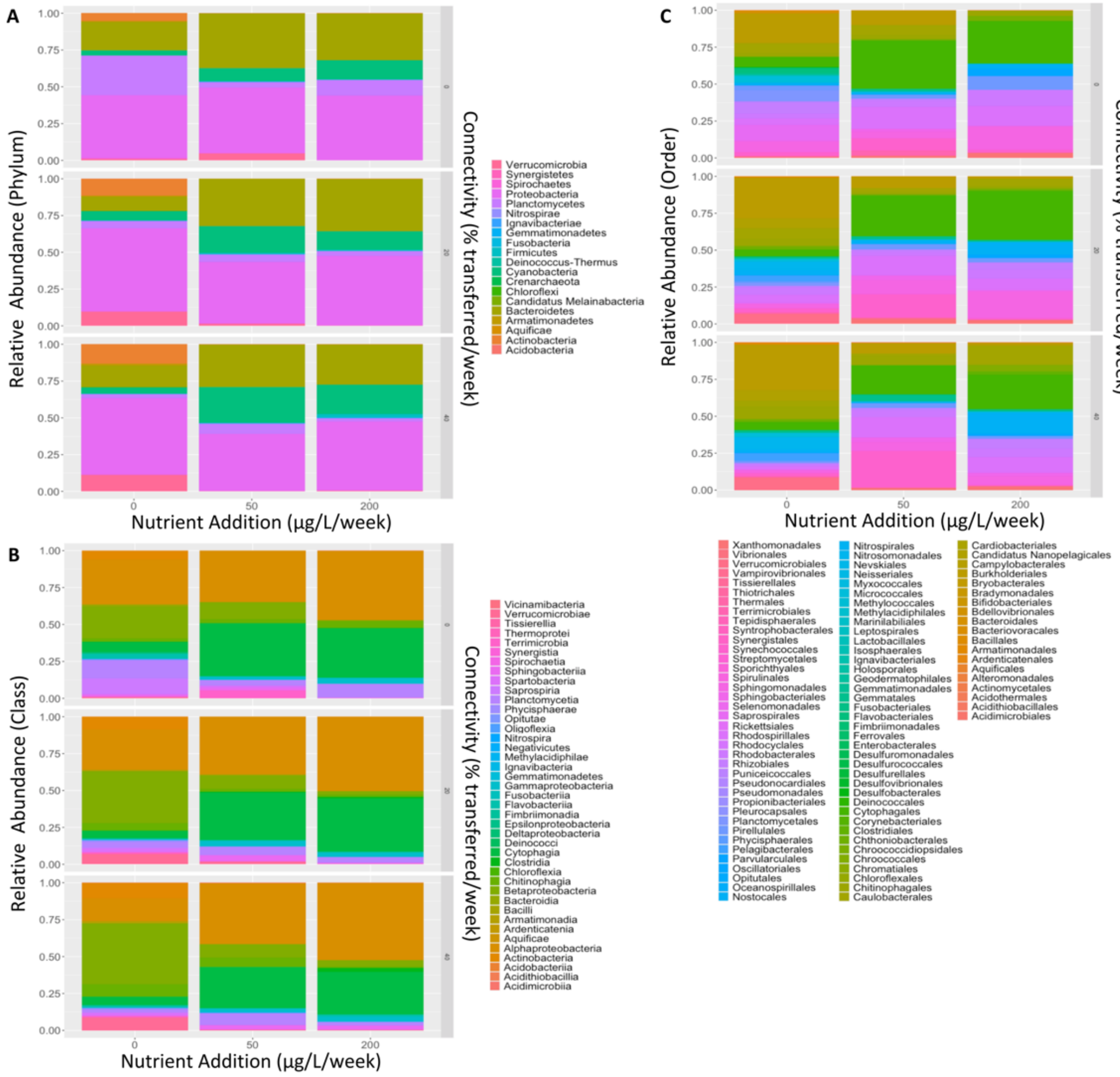


**Figure S1: Shifts in bacterioplankton community composition under the various combinations of nutrient enrichment and connectivity levels**. Shifts in community composition are visualized at the A) phylum, B) class, and C) order levels.

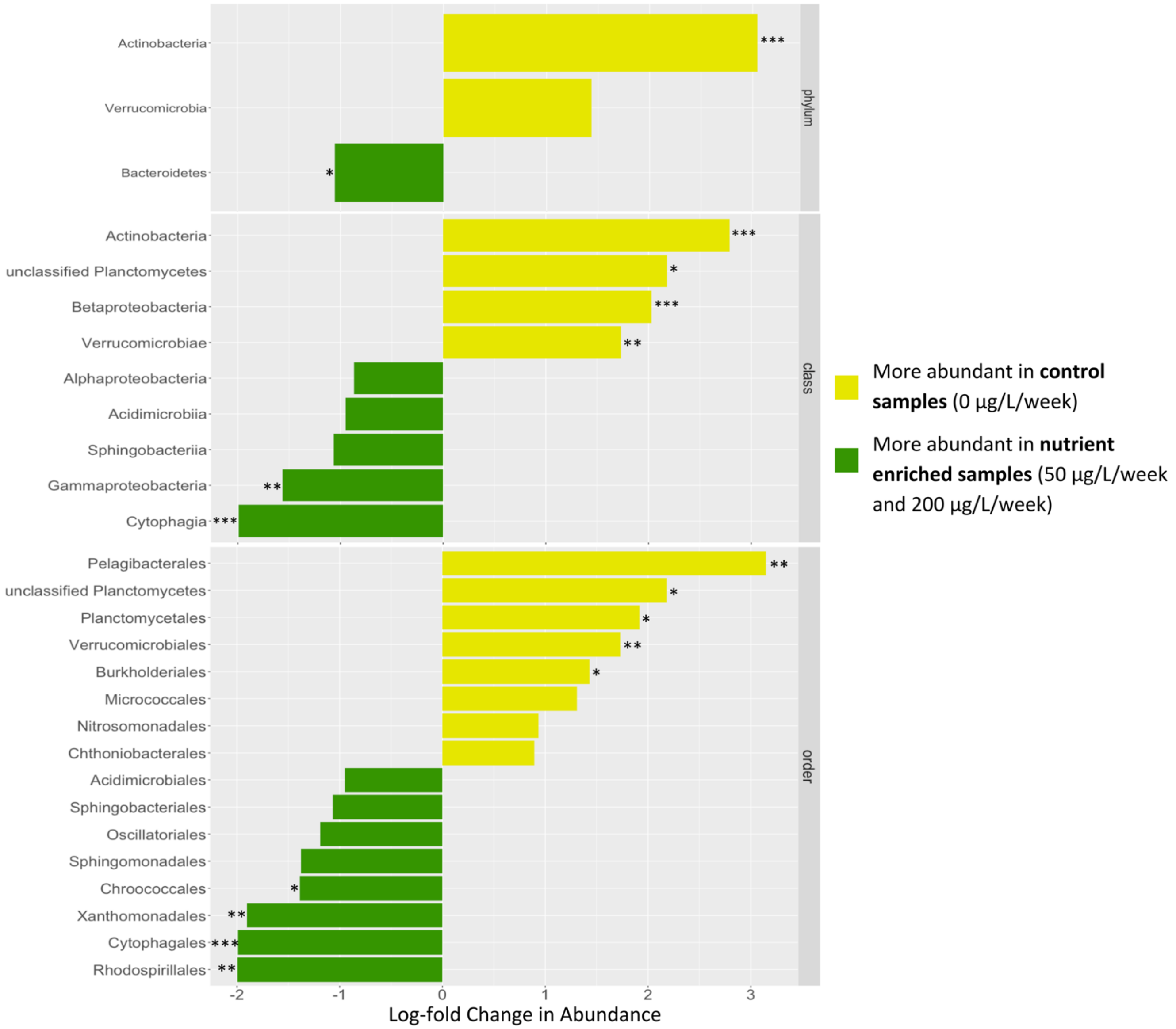


**Figure S2:** The results of the differential abundance analysis showing the differentially abundant taxa between the control (0 μg/L/week) and nutrient enriched (50 μg/L/week and 200 μg/L/week) samples at the phylum, class, and order levels. Only taxa present in at least 10 percent of samples were included in the analysis. Taxa with significant differences in abundance prior to multiple testing correction are shown here. (*) holm-adjusted $p < 0.05$, (**) holm-adjusted $p < 0.01$, (***) holm-adjusted $p < 0.001$

| observed species richness ~ connectivity + nutrient + connectivity:nutrient + sampling_date | | | | |
|---|---|---|---|---|
| | **Connectivity** | **Nutrient** | **Connectivity:Nutrient** | **Sampling Date** |
| **Estimate** | 2.820e-03 | -3.734e-03 | 6.061e-05 | -2.638e-01 |
| **Standard Error** | 1.441e-03 | 3.786e-04 | 1.352e-05 | 3.586e-02 |
| **Z-value** | 1.961 | -9.863 | 4.483 | -7.355 |
| **P-value** | 0.0499* | < 2e-16 *** | 7.34e-06 *** | 11.90e-13 *** |

| shannon diversity ~ connectivity + nutrient + connectivity:nutrient + sampling_date | | | | |
|---|---|---|---|---|
| | **Connectivity** | **Nutrient** | **Connectivity:Nutrient** | **Sampling Date** |
| **Estimate** | 8.837e-03 | -4.656e-03 | 4.778e-05 | -3.048e-01 |
| **Standard Error** | 7.226e-03 | 1.497e-03 | 5.841e-05 | 1.656e-01 |
| **T-value** | 1.223 | -3.110 | 0.818 | -1.841 |
| **P-value** | 0.23122 | 0.00418 ** | 0.41998 | 0.07585 |

**Table S4:** Results of the generalized linear model for both species richness fit to a Poisson distribution and Shannon diversity fit to a Gaussian distribution. Intercept for species richness model: estimate = 4.585e+00, standard error = 4.470e-02, z-value = 102.579, p-value = < 2e-16. Intercept for Shannon diversity model: estimate = 3.256e+00, standard error = 2.442e-01, t-value = 13.335, p-value = 6.7e-14.

| observed functional richness ~ connectivity + nutrient + connectivity:nutrient + sampling_date | | | | |
|---|---|---|---|---|
| | **Connectivity** | **Nutrient** | **Connectivity:Nutrient** | **Sampling Date** |
| **Estimate** | -9.257e-04 | -2.152e-03 | 7.259e-05 | -1.786e-01 |
| **Standard Error** | 2.878e-03 | 6.630e-04 | 2.422e-05 | 6.742e-02 |
| **Z-value** | -0.322 | -3.245 | 2.998 | -2.649 |
| **P-value** | 0.74770 | 0.00117 ** | 0.00272 ** | 0.00807 ** |

**Table S5:** Results of the generalized linear model for functional richness fit to a Poisson distribution. Intercept for species richness model: estimate = 3.400e+00, standard error = 7.886e-02, z-value = 43.114, p-value = < 2e-16.

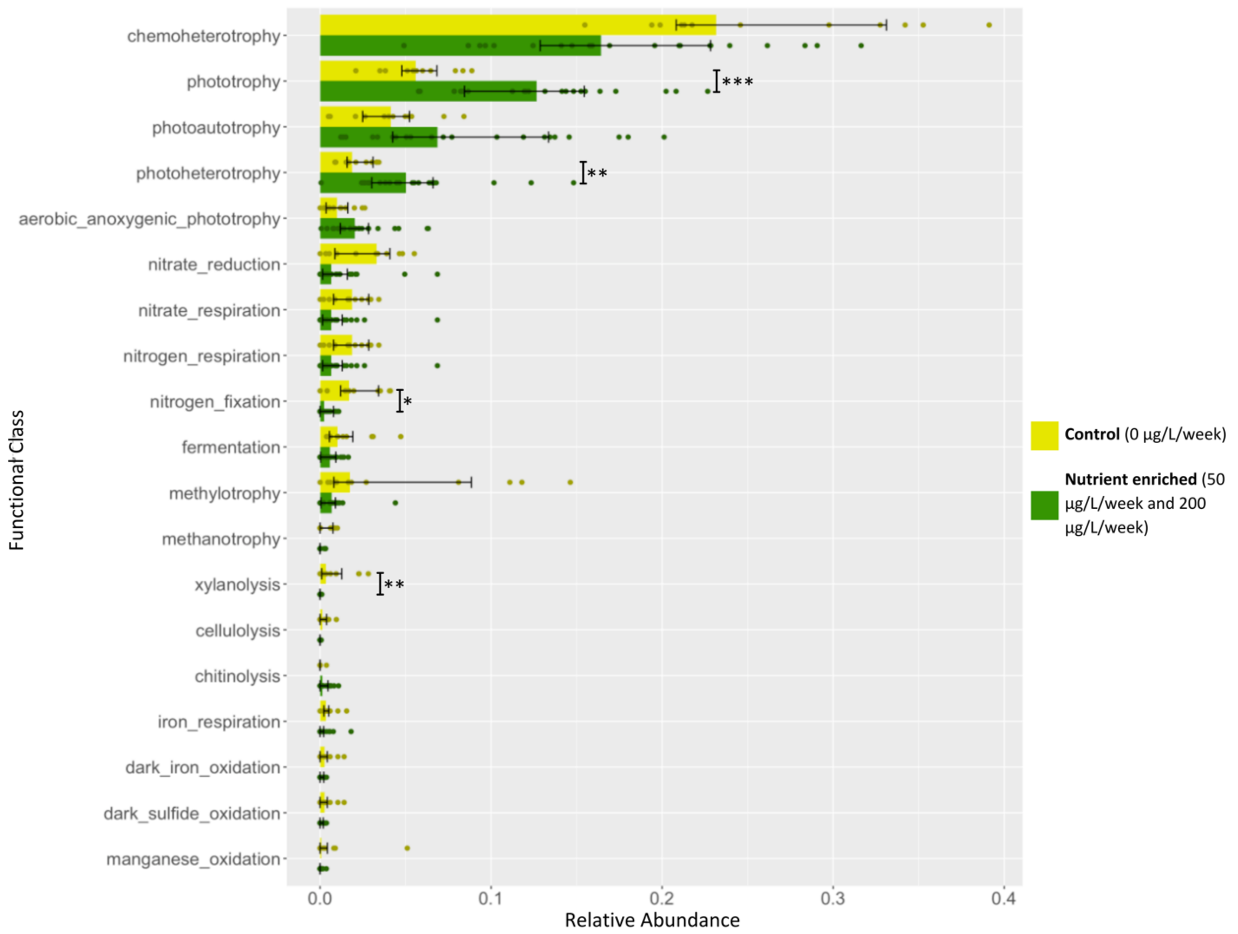


**Figure S3: Shifts in the abundance of functional classes between the control (0 µg/L/week) and nutrient enriched (50 µg/L/week and 200 µg/L/week) samples.** Only functional classes with significant differences in abundance prior to multiple testing correction are shown here. Asterisks indicated significant differences after multiple testing correction: (*) $p < 0.05$, (**) $p < 0.01$, (***) $p < 0.001$
